# *Escherichia coli* small heat shock protein IbpA is an aggregation-sensor that self-regulates its own expression at a translational level

**DOI:** 10.1101/2020.04.20.050740

**Authors:** Tsukumi Miwa, Yuhei Chadani, Hideki Taguchi

**Author notes:** Corresponding author (HT).

## Abstract

Aggregation is an inherent characteristic of proteins. Risk management strategies to reduce aggregation are critical for cells to survive upon stresses that induce aggregation. Cells cope with protein aggregation by utilizing a variety of chaperones, as exemplified by heat-shock proteins (Hsps). The heat stress-induced expression of IbpA and IbpB, small Hsps in *Escherichia coli*, is regulated by the σ^32^ heat-shock transcriptional regulator and the temperature-dependent translational regulation via mRNA heat fluctuation. We found that, even without heat stress, either the expression of aggregation-prone proteins or the *ibpA* gene deletion profoundly increases the expression of IbpA. Combined with other evidence, we propose novel mechanisms for the regulation of the small Hsp expression. Oligomeric IbpA self-represses the *ibpA/ibpB* expression at the translational level, but the self-repression is relieved by the sequestration of IbpA into protein aggregates. Thus, the function of IbpA as a chaperone to form co-aggregates is harnessed as an aggregation sensor to tightly regulate the IbpA level. Since the excessive preemptive supply of IbpA in advance of stress is harmful, the prodigious and rapid expression of IbpA/IbpB on demand is necessary for IbpA to function as a first line of defense against acute protein aggregation.

**Author summary:** All organisms have protein quality control systems against stresses disturbing cellular protein homeostasis (proteostasis). The systems have multiple stages: folding, degradation, and sequestration. Sequestration of denatured proteins is the first step to support other maintenance strategies. Small heat shock proteins (sHsps), which are well-conserved chaperones, are representative “sequestrases” that co-aggregate with denatured proteins. We found that IbpA, an *Escherichia coli* sHsp, is a direct mediator for negative feedback regulation at the translational level. Recruitment of IbpA into the protein aggregates relieves the *ibpA* expression suppression. This novel mechanism of IbpA as an aggregation-sensor tightly regulates the IbpA level, enabling the sHsp to function as a sequestrase upon aggregation stress.

## Introduction

Since proteins tend to form aggregates, cellular maintenance by keeping proteins in their native states and removing denatured proteins is crucial for all organisms. Multi-layered quality control systems are essential to maintain such cellular protein homeostasis (proteostasis) [1,2]. Refolding and degradation of denatured proteins caused by stresses are two primary strategies to prevent the accumulation of protein aggregates. Sequestration of denatured proteins is a third strategy, to keep misfolded proteins in a state that is easy to restore or degrade after stresses [1,2]. Small heat shock proteins (sHsps) participate in the third strategy as “sequestrases”, constituting a first line of stress defense against irreversible protein aggregation [3–6].

sHsps, defined as having low subunit molecular weights (12-43 kDa), are widely conserved chaperones from bacteria to mammals [3–6]. sHsps protect denatured proteins from forming irreversible aggregates by co-aggregating with the denatured proteins, in an ATP-independent manner [1,3–6]. The denatured proteins co-aggregated with sHsps can then be efficiently processed by other chaperones [1,3–6]. Although the minimum physiological unit of sHSPs is a dimer, sHsps usually form various types of oligomers, which are required for the energy-independent sequestration activity [4–6]. sHsps are composed of three domains, a flexible N-terminal domain, a highly conserved α-crystallin domain, and a C-terminal domain with an oligomerization motif, containing the characteristic three-residue IX(I/V) motif [4–6]. The C-terminal IX(I/V) motif functions as a cross-linker for intermolecular binding among dimeric sHsps [4–6].

sHsps are among the most upregulated Hsps upon stress [4,6,7]. The gene expression of the α- and γ-proteobacterial sHsps is regulated by two mechanisms [7]. One is the heat-shock transcriptional regulator σ^32^, an RNA polymerase subunit, which regulates the transcription of many Hsp genes [8,9]. At normal growth temperatures, σ^32^, which is an extremely unstable protein, is rapidly degraded via a DnaK/DnaJ-mediated pathway. However, σ^32^ is stabilized to allow the transcription of many Hsp genes upon heat shock, since DnaK/J is sequestered to rescue the emerging heat-denatured proteins [8,9]. The other is the thermoresponsive mRNA structures in the 5’ untranslated region (UTR), called RNA thermometers (RNATs), which mask the Shine-Dalgarno (SD) sequence in their stem loop structures at normal or low temperatures [7,10]. The heat fluctuation by a temperature up-shift melts the stem loops in RNATs and allows the ribosome to initiate translation, using the exposed SD sequence [7,10]. The thermo-responsivity of many bacterial RNATs has been established, and the RNATs of sHsps have conserved shapes harboring two to four stem loops [7,10]. Thus, the expression of sHsps is controlled at both the transcriptional level using heat-shock transcriptional factors, and the translational level using RNATs, in contrast with other Hsps, which are only controlled at transcriptional levels [7,10].

Inclusion body-associated protein A (IbpA) and B (IbpB) are *Escherichia coli* sHsps, and were originally identified as proteins induced in response to heterologous protein expression in the cell [11,12]. IbpA/IbpB function as a holder of denatured proteins, to facilitate the initiation step of the refolding pathway via DnaK/DnaJ and ClpB [4–6]. IbpA/IbpB mediate the efficient transfer of denatured proteins from the sHsp co-aggregation to the DnaK/DnaJ system [13–16]. IbpA and IbpB are highly homologous proteins with ~50% amino acid sequence identity [11], and form hetero-oligomers in *E. coli* [14,17,18]. Although most γ-proteobacteria only have a single sHsp (IbpA), a two-sHsp (IbpA and IbpB) system has evolved in a subset of *Enterobacterales*, including *E. coli*, from a single gene duplication event [19]. Previous *in vitro* studies using recombinant IbpA and IbpB proteins revealed the distinct features of the two sHsps: IbpA is more efficient in binding denatured proteins to form the coaggregates [17,19], and even self-forms fibril-like aggregates [18]. However, the coaggregates with IbpA are inefficient substrates for the disaggregation assisted by DnaK/DnaJ and ClpB [17,19]. The additional presence of IbpB in the coaggregates is required for the disaggregation process, suggesting that the interplay between IbpA and IbpB is important to modulate the interactions with substrates in the mixed oligomer states [4,6,16,17,19].

The mRNA encoding the *ibpA-ibpB* operon has RNATs in the 5’ UTRs of both the *ibpA* and *ibpB* ORFs, as revealed by RNA structure probing and reporter assays [7,20,21]. Previous analyses of the RNAT in *ibpB* using a reporter revealed the possible influence of the IbpA protein on *ibpB* expression [7,21]. In addition to the heat stress, the expression level of IbpA/IbpB was profoundly upregulated, by 10~50-fold, under non-heat stressed conditions such as 30~37 °C in the *dnaK-dnaJ* deleted strain [22,23] or upon oxidative stress induced by copper [24]. Since the RNAT regulation would not be effective at normal growth temperatures, the mechanism for the massive upregulation under the non-heat stressed conditions remains to be elucidated.

Here we addressed why IbpA is upregulated in non-heat stressed cells. We found that the accumulation of protein aggregates was sufficient for the upregulation. Intriguingly, a reporter assay using the 5’ UTR of the *ibpA* mRNA revealed that the deletion of the *ibpA* gene increased the reporter translation, which was repressed by the overexpression of oligomeric IbpA. Combined with other evidence, we propose that the IbpA oligomers self-repress the *ibpA* translation, which is relieved by the sequestration of IbpA by co-aggregation with protein aggregates. The role of the aggregation sensor is specific to IbpA, since the homologous IbpB lacks this self-repression function. The significance of the self-repression by IbpA at the translational level is discussed in relation to the unique role of IbpA in protecting cells from acute heat stress.

## Results

### *ibpA* translation is upregulated in response to protein aggregation

Although the thermometer in the mRNA (RNAT) and the transcriptional control by σ^32^ are known mechanisms to upregulate the expression of IbpA, previous studies have reported that IbpA expression is also upregulated under non-heat stressed conditions, such as in the *dnaKJ* deletion strain or upon copper stress [22–24]. The absence of DnaK/DnaJ leads to the production of protein aggregates [22,25]. The addition of copper disturbs protein homeostasis in cells with oxidative stress [24,26]. A common consequence in *E. coli* cells would be the accumulation of protein aggregates [20–23]. Therefore, we hypothesized that protein aggregation might somehow be involved in the upregulation of IbpA under the non-heat stressed conditions.

To investigate whether the expression of IbpA is upregulated not only by heat shock but also by protein aggregation under non-heat stress conditions, we expressed aggregation-prone proteins in wild-type *E. coli*. To do so, we overexpressed rhodanese, a bovine mitochondrial protein, which is known to aggregate in *E. coli* at 37 °C [27] (S1*A* Fig.). Strikingly, the expression of IbpA increased upon the rhodanese expression (*agg*^++^, Fig. 1*A*). Overexpression of another aggregation-prone protein, SerA of *E. coli* [25] (S1*A* Fig.), also massively induced the IbpA expression in wild-type *E. coli* (S1*B* Fig.). The rhodanese expression did not increase the level of GroEL, one of the representative Hsps in *E. coli* (Fig. 1*A*). These results support the idea that the accumulation of aggregated proteins in cells increases the IbpA expression.

**Fig 1.**
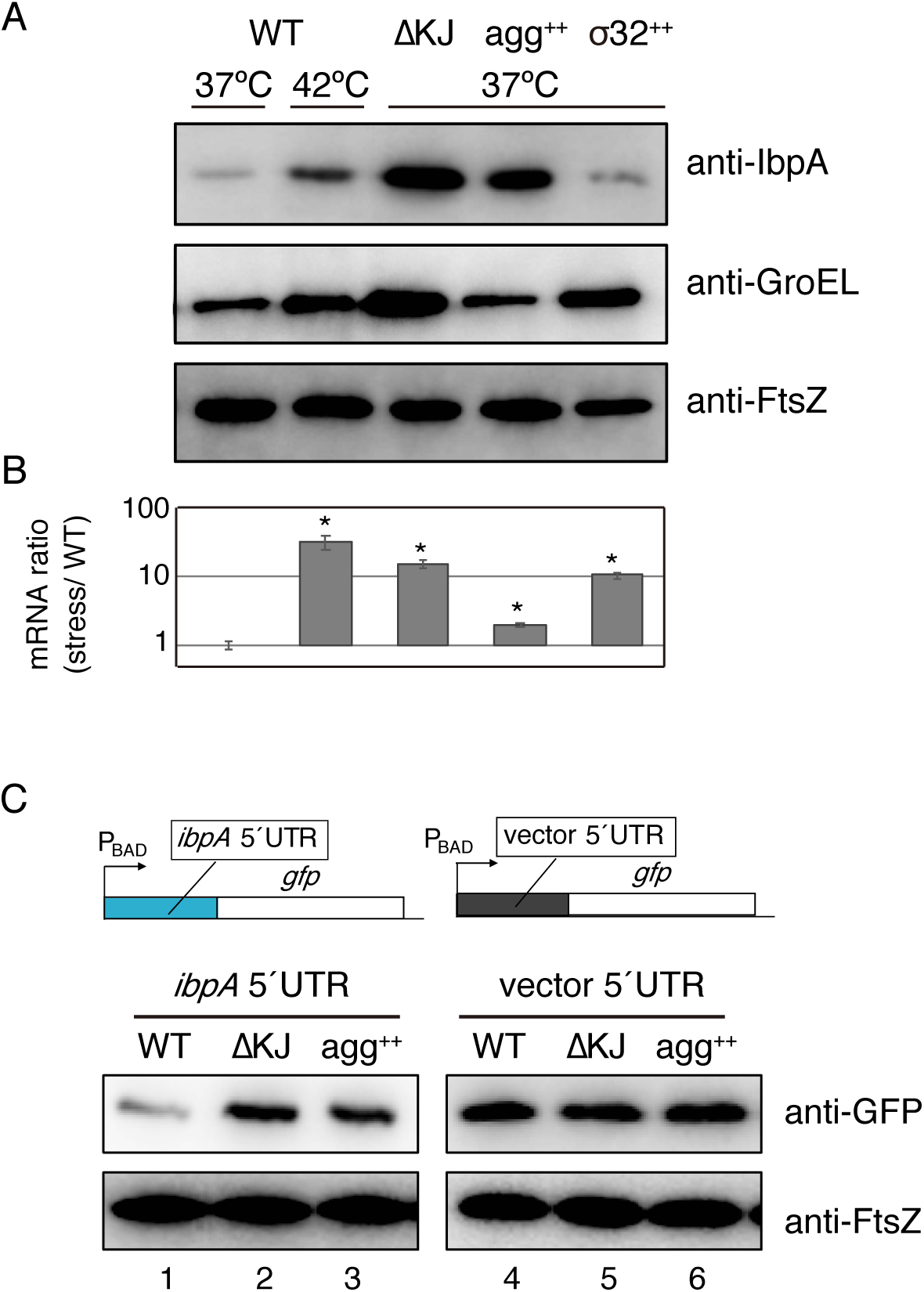
Accumulation of protein aggregates upregulates *ibpA* translation at the translational level. (*A*) Western blotting to evaluate the endogenous *ibpA* expression in *E. coli* wild-type (*WT*) cells under various conditions. *E. coli* cells were grown at 37 °C or shifted to 42 °C for 10 min. ∆*KJ*, the *dnaKJ* deletion strain; *agg*^++^, *E. coli* wild-type expressing rhodanese; σ^32++^, *E. coli* wild-type strain expressing σ^32^. Unless otherwise indicated, *E. coli* cells were grown at 37 °C. Expressions of GroEL and FtsZ were also examined as controls for a typical Hsp and a constitutively expressed protein, respectively. (*B*) Ratios of the *ibpA* mRNA amounts in cells under conditions corresponding to (*A*). Error bars represent SD; *n* =3 biological replicates. Student’s *t* test was used to assess the statistical significance of differences (*: *p* < 0.01). (*C*) Evaluation of *ibpA* translation by GFP reporters. The reporters harboring the 5’ UTR of *ibpA* or the 5’ UTR from a plasmid without the *ibpA* sequence were expressed under various conditions. Western blotting was performed using anti-GFP and anti-FtsZ antibodies.

We suspected that the aggregates might induce IbpA via upregulated σ^32^-mediated transcriptional control at 37 °C, since the aggregation-prone proteins could sequester DnaK/J, thus protecting σ^32^ from DnaK/J-mediated degradation and eventually stabilizing σ^32^ to promote the expression of Hsps. If so, then the overexpression of σ^32^ would increase the mRNAs to upregulate IbpA as well as other chaperones, such as GroEL. After we confirmed the ~10-fold induction of *ibpA* mRNAs in the σ^32^-overexpressing cells, using quantitative real-time PCR (qRT-PCR) (Fig. 1*B*), we compared the protein expression levels. Upon the σ^32^ overexpression, the GroEL expression level increased (Fig. 1>*A*), indicating that the excess σ^32^ is effective in increasing the Hsp under the non-heat stress. In contrast, the expression level of IbpA did not obviously increase as compared to that of GroEL (Fig. 1*A*), even though the *ibpA* mRNA increased, indicating that the IbpA induction in the presence of aggregates is not explained by transcriptional upregulation using σ^32^. Rather, the results suggest a translation suppression mechanism in the IbpA expression.

To demonstrate that the *ibpA* upregulation by the accumulation of protein aggregates does not occur at the transcriptional level, we investigated the efficiency of the *ibpA* translational initiation in a reporter assay. Since the 5’ UTR in the *ibpA* mRNA is critical for the translational control using the stem loops in the mRNA, we constructed a plasmid harboring the 5’ UTR of *ibpA* fused with the *gfp* gene, under the control of an arabinose-inducible promoter (Fig. 1*C*). The deletion of *dnaK/J* (Δ*KJ*) caused a massive increase in the GFP production upon arabinose induction, verifying that the reporter reflected the features of the IbpA upregulation (Fig. 1*C*). We observed a similar substantial induction of GFP upon the rhodanese overexpression. The effect is specific for the 5’ UTR of *ibpA*, since there was no upregulation of GFP in the ∆*KJ* cells or upon the rhodanese overexpression when the 5’ UTR was substituted with a 5’ UTR derived from the parent plasmid. In addition, the overexpression of another aggregation-prone protein, SerA, also induced the GFP reporter (S1*B* Fig.). The results using the reporter, where the transcriptional levels controlled by the arabinose-inducible promoter were independent of heat-shock, further confirmed that the accumulation of protein aggregates upregulates the *ibpA* expression at the translational level.

### IbpA self-represses *ibpA* translation

Next, we investigated the connection between protein aggregation and *ibpA* translation induction. The well-known physiological function of IbpA as a chaperone is the co-aggregation with denatured proteins [1,3–6]. The recruitment of IbpA to protein aggregates might cause the entrapment of the free IbpA in the cytosol. Indeed, the localization of IbpA fused with GFP shifted to the cell poles in the DnaK/J-depleted cells and the rhodanese-overexpressing cells (Fig. 2*A*), consistent with previous observations that IbpA and denatured proteins accumulate as inclusions at the cell poles [28–30]. Taking this into consideration, we hypothesized that the entrapment of free IbpA from the cytosol, due to the sequestration of denatured proteins, would induce the *ibpA* translation (Fig. 2*B*). If this model is correct, then the deletion of IbpA, which is nonessential for *E. coli* growth, would upregulate the translation of the reporter harboring the 5’ UTR of *ibpA*, used in Fig. 1*C*.

**Figure 2.**
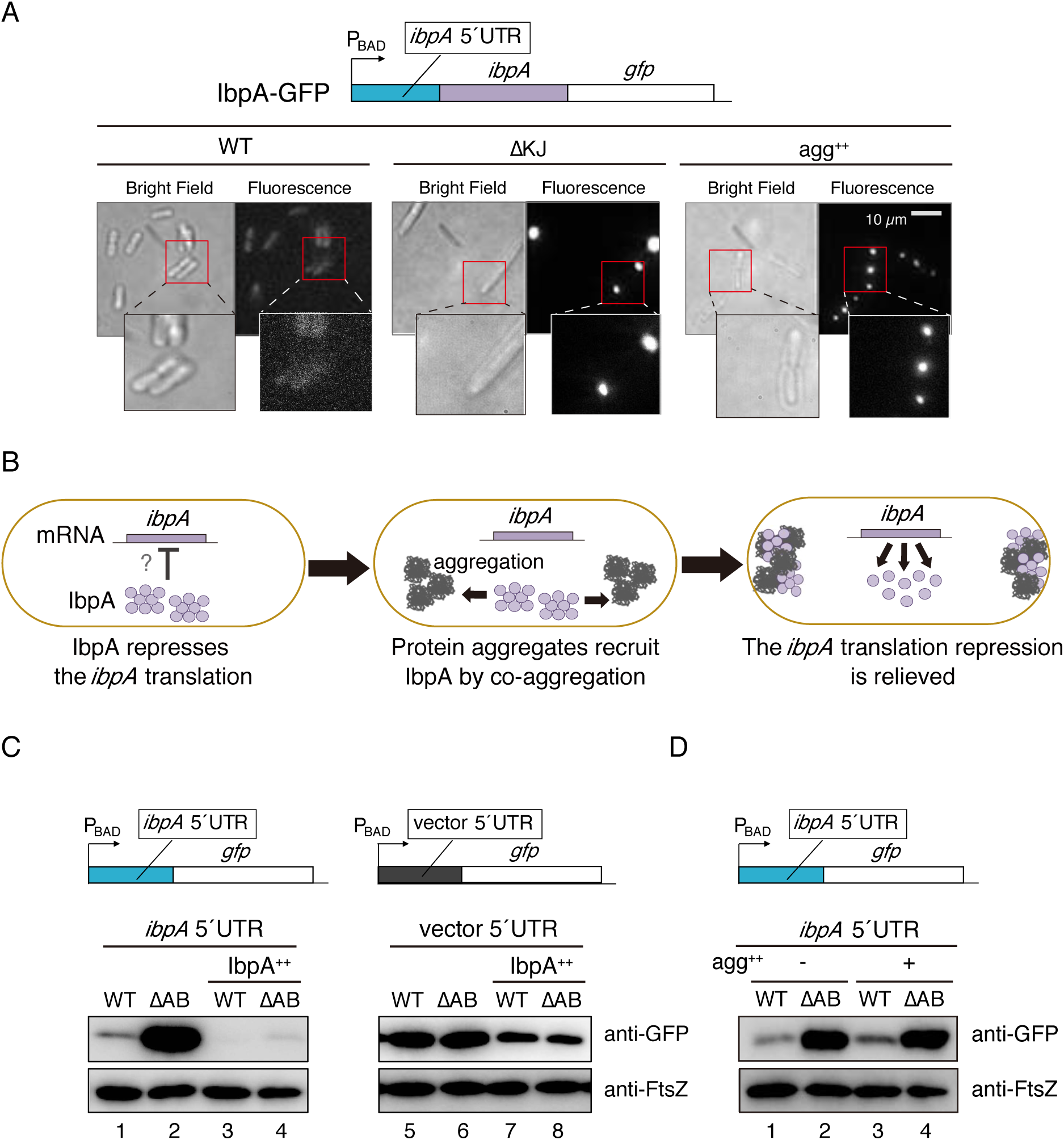
IbpA self-represses *ibpA* translation. (*A*) Localization of IbpA-GFP in *E. coli* cells. *Top*. The IbpA-GFP reporter construct is schematically shown. *Bottom*. Bright-field and fluorescence images of IbpA-GFP in the absence or presence of induced protein aggregation. *WT*, *E. coli* wild-type strain; ∆*KJ*, the *dnaKJ* deletion strain; *agg*^++^, wild-type strain expressing rhodanese. (*B*) Titration model of the IbpA-mediated negative feedback mechanism. IbpA (*circles*) suppresses its own mRNA translation. Upon aggregate (*dark blobs*) formation, the IbpA co-aggregated with aggregation-prone proteins is sequestered at the cell poles, relieving the translation suppression. (*C*) Evaluation of the *ibpA* translation initiation by the reporters used in Fig. 1*C*. *WT*, *E. coli* wild-type strain; ∆*AB*: the *ibpAB* operon deleted strain; *IbpA*^++^, IbpA was induced by 0.1 mM IPTG for 1 h. Western blotting with anti-GFP or anti-FtsZ antibodies is shown. (*D*) Evaluation of the *ibpA* translation initiation by the GFP reporter used in Fig. 1*C*. *E. coli* lysates from wild-type (*WT*) or the *ibpAB* operon-deleted strain (∆*AB*) with (+) or without (–) the co-expression of rhodanese (*agg*^++^) were analyzed. Western blotting using anti-GFP and anti-FtsZ antibodies is shown.

We deleted the operon including *ibpA*-*ibpB*. After we confirmed that the growth of the *∆ibpAB* cells was similar to that of wild-type *E. coli* (S2*A* Fig.), we evaluated the translation initiation of the reporter. Strikingly, the reporter expression was strongly promoted by the *ibpAB* deletion, even in the absence of induced protein aggregation (Fig. 2*C*, *lane 2*), supporting the idea that IbpA suppresses *ibpA* translation. We then examined the effect of IbpA overexpression, which caused a slower growth rate in wild-type *E. coli* (S2*B* Fig.). The overexpression of IbpA in wild-type cells completely suppressed the expression of the reporter (Fig. 2*C*, *lane 3*). More importantly, the replenishment of IbpA by the overexpression in the ∆*ibpAB* cells almost completely repressed the upregulated expression of the GFP reporter (Fig. 2*C*, *lane 4*). Further rhodanese overexpression did not increase the upregulated expression level of the reporter in the ∆*ibpAB* cells (Fig. 2*D*). The lack of additional effects by the aggregates on the IbpA induction suggests that the abundance of IbpA in cells governs the *ibpA* translation.

### Oligomeric IbpA is critical for the self-repression of translation

What region of IbpA is critical for the self-regulation of translation? At first we deleted the N- or C-terminal domain of IbpA and examined the effect on translation repression (Fig. 3*A*). The GFP reporter assay revealed that the C-terminal truncation eliminated the ability to suppress the translation in the ∆*ibpAB* cells (Fig. 3*A*). In contrast, the translation suppression by the N-terminal truncation was almost the same as that by wild-type IbpA (Fig. 3*A*), showing that the C-terminal domain is responsible for the self-translation repression.

**Figure 3.**
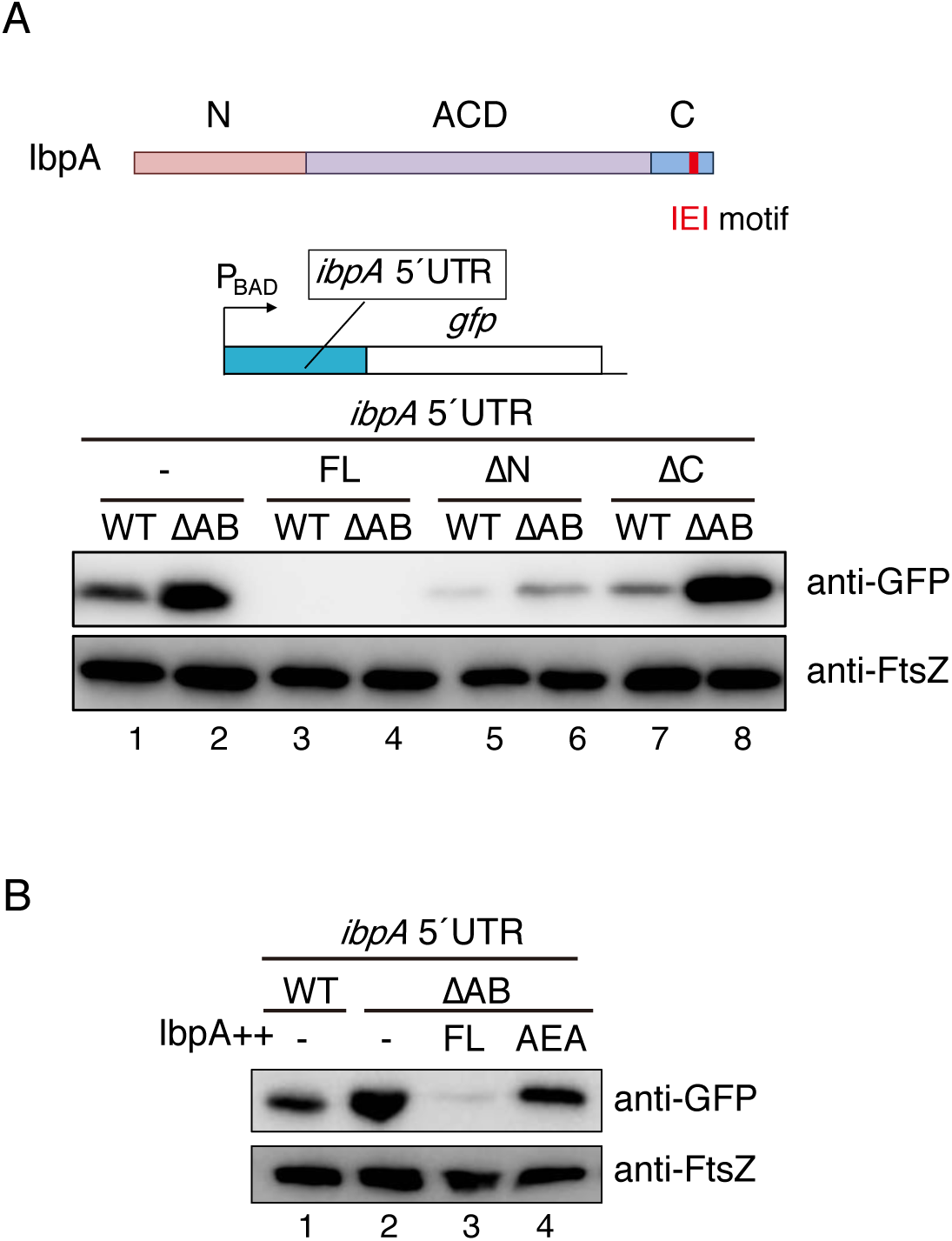
IbpA oligomerization is critical for the self-regulation. (*A*) Schematic representation of IbpA domains. *N*, N-terminal domain; *ACD*, α-crystallin domain; *C*, C-terminal domain. *Red*: the IEI motif, the IX(I/V) motif in IbpA. (*B*) The GFP reporter for the *ibpA* translation initiation used in Fig. 1*C* was expressed in wild-type *E. coli* (*WT*) or the *ibpAB*-deleted strain (∆*AB*) co-expressing the full-length IbpA (*FL*), N-terminal domain-deleted mutant (∆*N*), and C-terminal domain-deleted mutant (∆*C*). (*C*). In addition to wild-type IbpA (FL), the IbpA(AEA) mutant (*AEA*) was evaluated as in (*B*). Western blotting with anti-GFP or anti-FtsZ antibodies is shown.

The C-terminal domain of IbpA contains a universally conserved motif for oligomerization, IX(I/V) [31,32]. We substituted the motif, IEI in IbpA, with AEA, and confirmed the impaired oligomerization of the AEA mutant (S3 Fig.), as reported previously [31]. Co-expression of the IbpA (AEA) mutant with the reporter in the ∆*ibpAB* cells did not repress the *ibpA* translation (Fig. 3*B*), indicating that the oligomeric state of IbpA is critical for the IbpA self-regulation.

### IbpA also suppresses the translation of *ibpB*

*E. coli* also possesses IbpB, a paralog of IbpA, and its expression is also regulated by RNAT in the 5’ UTR of the *ibpB* mRNA [7,20,21]. Gaubig *et al.* observed that the presence of the *ibpA* ORF suppresses the *ibpB* translation [21], but the reporter system used in the study could not reveal the influence of IbpB on the *ibpB* translation. Therefore, we tested the effect of IbpB on the *ibpB* translation, after we replaced the 5’ UTR of *ibpA* with that of *ibpB* in the GFP reporter system. IbpA, but not IbpB, suppressed the reporter for the *ibpB* translation in the ∆*ibpAB* strain (Fig. 4*A*). This result shows the specific function of IbpA as a translation repressor of the small Hsps, IbpA and IbpB, in *E. coli*.

**Figure 4.**
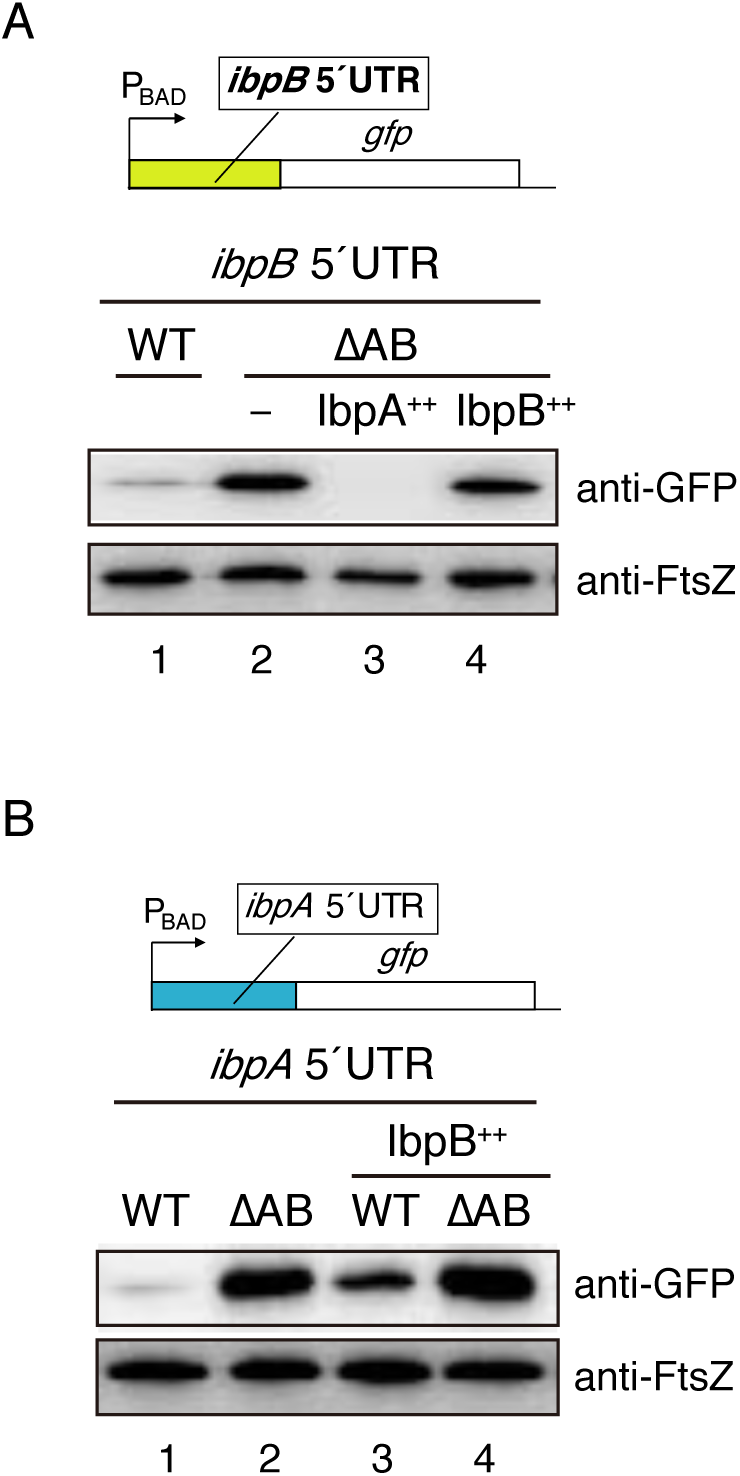
IbpB cannot substitute for IbpA as the sHsp translation suppressor. (*A*) Evaluation of the *ibpB* translation initiation by IbpA or IbpB. The 5’ UTR of *ibpA* in the GFP reporter used in Fig. 1*C* was replaced with the 5’ UTR of *ibpB*. The modified GFP reporter was expressed in wild-type *E. coli* (*WT*) or the *ibpAB*-deleted strain (∆*AB*) co-expressing either IbpA or IbpB. (*B*) Effect of IbpB overexpression on the *ibpA* translation initiation evaluated by the GFP reporter used in Fig. 1*C*. Western blotting with anti-GFP and anti-FtsZ antibodies is shown.

In contrast, the overexpression of IbpB did not change the *ibpA* reporter upregulation in the ∆*ibpAB* strain (Fig. 4*B*, *lane 4*), indicating that IbpB cannot substitute for IbpA in the self-repression of the *ibpA* translation. In the wild-type strain, the IbpB overexpression increased the amount of the GFP reporter (Fig. 4*B*, *lane 3*), probably reflecting the hetero-oligomerization of IbpB with endogenous IbpA to reduce the amount of free IbpA for the self-repression.

### Stem loops in the 5’ UTR of *ibpA* mRNA mediate the self-repression of *ibpA* translation

Previous studies revealed that the secondary mRNA structures of the *ibpA* 5’ UTR regulate the *ibpA* translation [20]. The *ibpA* 5’ UTR contains three stem loops, and the two upstream stem loops (SL1 and SL2) are thought to stabilize the downstream thermo-responsive stem loop (SL3) to mask the SD sequence (Fig. 5) or protect the mRNA from degradation [10,20]. Since the contribution of these stem loops to translational regulation remains unclear, we constructed a series of stem loop variants in which the stem loop stabilities were weakened or strengthened (S4 Fig.), and evaluated the effects on the GFP reporter. For all three weakened stem loops, the GFP expressions were upregulated in wild-type *E. coli* as compared to those of the unchanged stem loops (Fig. 5*A-C*, compare *lanes 1 and 4*). The upregulation levels in wild-type *E. coli* were almost the same as those in the ∆*ibpAB* cells (compare *lanes 4 and 5*), but were largely suppressed when IbpA was overexpressed (*lane 6*). These results suggest that the translation suppression by the endogenous level of IbpA was compromised in the reporters harboring the weak stem loops in SL1-3.

**Figure 5.**
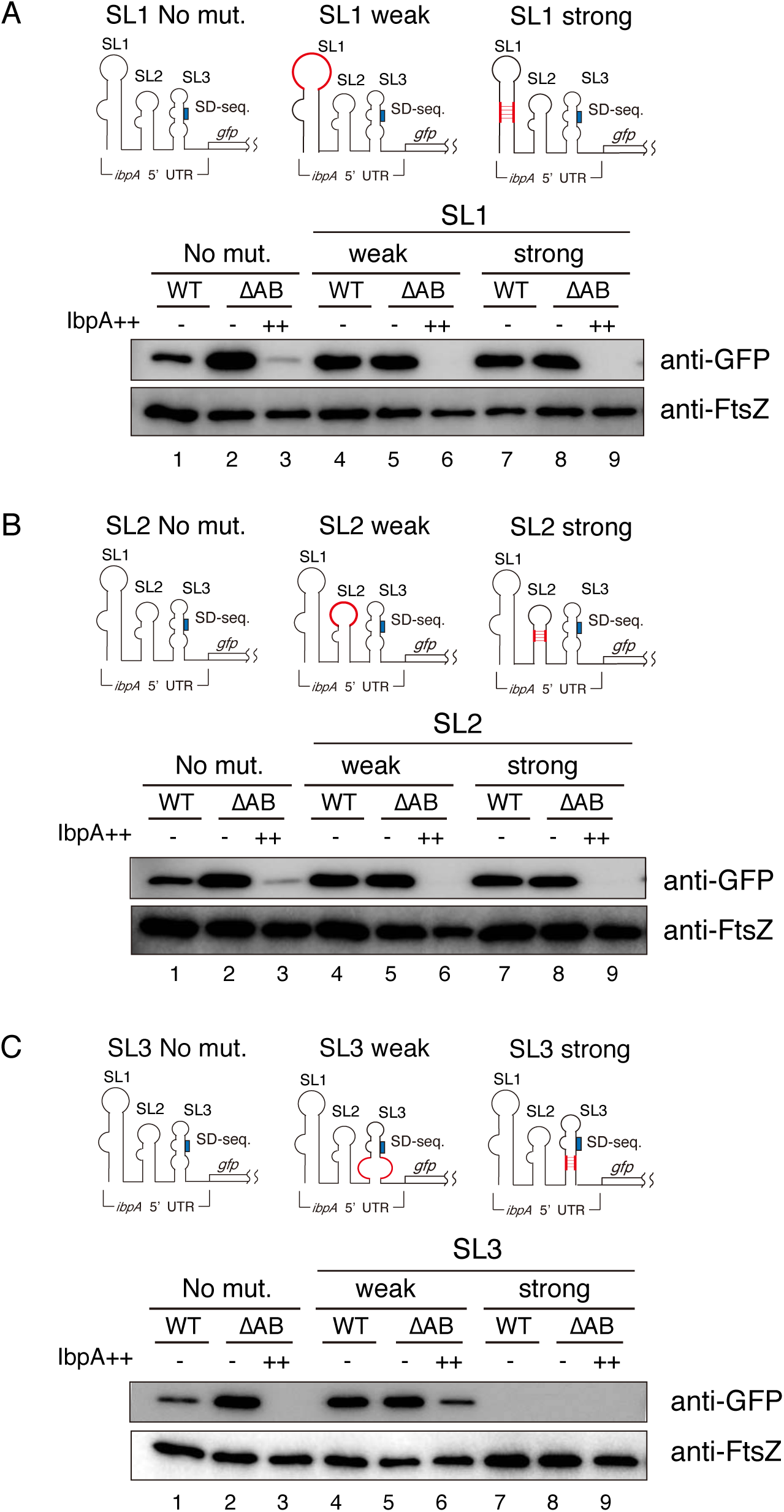
Effect of stem loops in the *ibpA* mRNA on the IbpA-mediated translation suppression. The stem loops in the *ibpA* 5’ UTR of the GFP reporter used in Fig. 1*C* were mutated and evaluated in the wild-type (*WT*) and ∆*ibpAB* (∆*AB*) strains. Mutations to weaken or strengthen the stem loop structures were introduced in SL1 (*A*), SL2 and SL3 (*C*). Schematic representations of stem loop mutations are shown (*see also SI Appendix*, Fig. S4). Where indicated, IbpA was overexpressed (++). The *ibpA* 5’ UTRs with no (*No mut*), weak and strong mutations in the stem loops were tested using the GFP reporter. Western blotting with anti-GFP and anti-FtsZ antibodies is shown.

The expression patterns of the reporters using the strong SL1 and SL2 stem loops were almost the same as those using the weak variants (Fig. 5*A, B*). In contrast, the expression of the reporter using the strong SL3 was not observed under all conditions examined (Fig. 5*C*), probably because the strong stem loop containing the SD motif is too tight to expose the SD motif for the translation initiation.

## Discussion

IbpA and IbpB, small Hsps in *E. coli*, are chaperones that sequester aggregation-prone proteins by co-aggregation during stress. This study unveiled a previously unknown function of IbpA, as a mediator for negative feedback regulation at the translational level. Our experiments revealed that the IbpA oligomers serve as their own translation suppressor. The titration of IbpA by co-aggregation with denatured proteins relieves the translation suppression.

We propose a novel mechanism of the *ibpA* regulation, where IbpA-mediated aggregation sensing self-regulates the *ibpA* translation (Fig. 6). Without protein aggregation, free IbpA oligomers suppress *ibpA* translation initiation. In contrast, under stress conditions involving protein aggregate accumulation, IbpA recruits aggregation-prone proteins to form co-aggregates. This titrates the free IbpA away to relieve the translation suppression, leading to the massive increase in IbpA expression to maintain cellular proteostasis.

**Figure 6.**
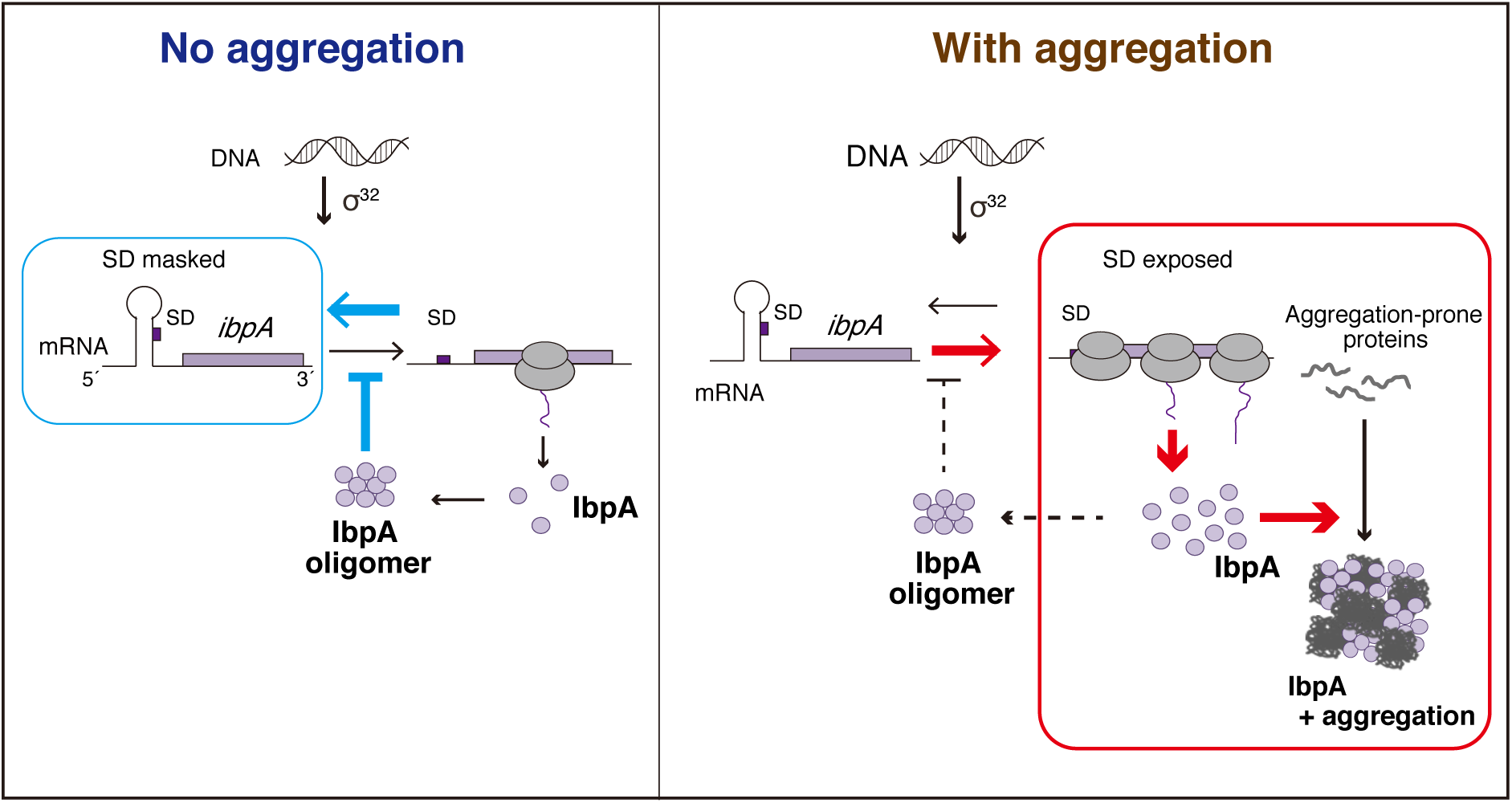
Model of the *ibpA* expression regulation. *Left*, Under normal conditions, the IbpA oligomers repress its own translation. *Right*, Under stress conditions with the accumulation of protein aggregates, the IbpA sequestration into protein aggregates relieves the IbpA-mediated translation suppression, leading to the prodigious expression of IbpA.

This model of IbpA expression regulation resembles the σ^32^-mediated transcriptional regulation of Hsps [8,9], since the chaperones are titrated away by denatured proteins under stress conditions in both mechanisms. One of the main differences between the two mechanisms is that the IbpA-mediated self-regulation takes place at the translational level, providing an advantage for a rapid response to the emergence of aggregation-prone proteins.

Previous studies revealed the layered regulation of IbpA expression: σ^32^-mediated transcriptional control and RNA thermometer (RNAT) translational control [7,21]. Since the stem loops in the RNAT system influence the IbpA-mediated translation repression (Fig. 5), RNAT and the self-repression control are not independent. In the RNAT mechanism, the stem loops fluctuate and melt to expose the SD region, depending on the temperature. As reported previously, higher temperatures cause more melting of the stem loops, like a “thermometer”. In other words, the degree of melting gradually changes, and is not all or none, over a wide range of temperatures [7,10]. This inherent property would allow the stem loops to partly open even under mild conditions such as 37 °C, leading to the leaky expression of certain amounts of IbpA. Thus, the IbpA-mediated self-repression would function to tightly shut off the IbpA expression as a “safety catch” in the leaky RNAT system. Taken together, the stringent repression mechanism, combining RNAT and the IbpA-mediated negative feedback control, has evolved to fulfill the following requirements: tight repression under unstressed conditions, and acute upregulation upon aggregation-stress. This mechanism enables IbpA to be one of the most upregulated chaperones upon aggregation inducing stresses.

IbpA serves as a first line of defense against protein aggregation, where oligomerized IbpA co-aggregates with aggregation-prone proteins for sequestration. Why does IbpA employ such a feedback control mechanism in addition to the known regulation controls including σ^32^ and RNATs? Considering that IbpA is an ATP-independent oligomeric chaperone, a greater than stoichiometric amount of IbpA would be necessary to sequester the aggregation-prone proteins. One strategy for risk management is to prepare an abundance of IbpA protein even under unstressed conditions. However, this might not be appropriate, since IbpA overexpression had detrimental effects under the normal conditions (S2*B* Fig.), probably due to the self-formation of fibril aggregates [18], which could perturb proteostasis and compromise the sequestration activity. Thus, the expression of IbpA should be tightly repressed under normal conditions, since IbpA can be regarded as a “double-edged sword”. The self-regulation mechanism proposed here can overcome the dilemma that the high abundance of IbpA is necessary in cases of aggregation stress, but an excessive preemptive supply could be detrimental to the cell.

Our analysis of the stem loops in the 5’ UTR of the *ibpA* mRNA revealed that the secondary structures of the mRNA are critical to regulate the translation. How does IbpA couple to the mRNA structure in RNAT for the translation suppression? One possibility is that the oligomeric states of IbpA bind to their own mRNA to suppress the translation, although no RNA-binding motif has been detected in IbpA. Our stem loop variants revealed that SL1/SL2 affect SL3 for the translation initiation, suggesting that SL1/SL2 interact with SL3 to control the translation. In this case, the IbpA oligomers might associate with the RNAT system to suppress the translation. Alternatively, a *trans*-acting modulator might contribute to the link between RNAT and IbpA.

*E. coli* has two small Hsps, IbpA and its highly homologous paralog IbpB, which are encoded in the *ibpAB* operon [11]. Several lines of evidence have shown the distinct roles of IbpA and IbpB, where IbpA and IbpB function as a canonical binder and its noncanonical paralog that enhances the dissociation of sHsps from the co-aggregates, respectively [17,19]. In addition to this distinction, our findings demonstrated another aspect of the difference between the two sHsps in the expression regulation. IbpA suppressed the *ibpB* reporter translation in the ∆*ibpAB* strain (Fig. 4, [21]). In contrast, IbpB could not suppress the *ibpA* translation. Thus, IbpA plays a pivotal role as a master regulator of the expression of sHsps at the translational level, ensuring that IbpA and IbpB cooperate to cope with protein aggregation.

The overexpression of IbpB in the wild-type strain increased the IbpA level (Fig. 4). This IbpA induction is interpreted to be due to the IbpA deprivation by hetero-oligomer formation between IbpA and IbpB, implying that, in addition to the aggregation-prone proteins, the factors that can associate with IbpA could trigger its upregulation. Therefore, we suggest the possibility that IbpA plays a pivotal role as a *trans*-regulator for the expression of other proteins. Indeed, the translation level of *ibpB* is decreased upon IbpA co-expression [21]. The fact that the *ibpB* mRNA also has an RNAT in the 5’ UTR [20,21] implies that IbpA recognizes a series of stem loops in 5’ UTRs, such as RNAT structures.

## Materials and methods

### *E. coli* strains

The *E. coli* DH5α strain was used for cloning. The BW25113 strain was used for each assay. Deletion of the chromosomal *ibpA-ibpB* operon or *dnaK-dnaJ* operon was accomplished by the procedures described previously [33]. The DNA fragment amplified from JW3664 (∆*ibpA*::FRT-Km-FRT), using the primers PT0456 and PM0195, and that from JW3663 (∆*ibpB*::FRT-Km-FRT), using the primers PT0457 and PM0196, were mutually annealed and amplified using PT0456 and PT0457. The purified DNA was electroporated into the *E. coli* strain BW25113 harboring pKD46, and the transformant resistant to 40 µg/ml kanamycin was stored as BW25113∆*ibpAB*. BW25113∆*dnaKJ* was constructed using JW0013 (∆*dnaK*::FRT-Km-FRT), JW0014 (∆*dnaJ*::FRT-Km-FRT), PT0071, PT0072, PM0195 and PM0196, as described above. Primers are listed in S1 Table.

### Plasmids

Plasmids were constructed using standard cloning procedures and Gibson assembly. Plasmids for reporter or microscopy assays: pBAD30-*ibpA* 5’ UTR-*gfp*, pBAD30-*gfp*, pBAD30-*ibpB* 5’ UTR-*gfp*, and pBAD30-*ibpA* 5’ UTR-*ibpA-gfp* were constructed using DNA fragments amplified from pBAD30 [34], DNA fragments amplified from superfolder GFP [35], derived from a plasmid constructed previously [36], and DNA fragments amplified from *E. coli* genomic DNA. Plasmids for overexpression: pCA24N-*rhodanese*, pCA24N-*rpoH*, pCA24N-*serA*, pCA24N-*gfp*, pCA24N-*ibpA*, pCA24N-*ibpA*_AEA and pCA24N-*ibpB* were constructed using DNA fragments amplified from pCA24N [37], DNA fragments amplified from superfolder GFP [35,36] or DNA fragments amplified from *E. coli* genomic DNA. Primers used for cloning are listed in S1 Table.

### SDS-PAGE and western blotting

*E. coli* BW25113 cells harboring a plasmid carrying the reporter were precultured at 30 °C for 16 h in LB medium. The precultured cells were grown to an OD_660_ of 0.4~0.6 at 37 °C in LB medium with 2 × 10^−4^ % arabinose. For the co-expression assay, *E. coli* BW25113 cells harboring plasmids carrying the reporter genes and pCA plasmids were used. The induction of protein co-expression was performed with 0.1 mM isopropyl-β-D-thiogalactopyranoside (IPTG), added 2 h after starting the culture. Cell cultures were sampled and mixed with an equal volume of 10% TCA, to stop the biological reactions and precipitate the macromolecules. After standing on ice for at least 15 min, the samples were centrifuged for 3 min at 4 °C, and the supernatant was removed by aspiration. Precipitates were washed with 1 mL of acetone by vigorous mixing, centrifuged again, and dissolved in 1× SDS sample buffer (62.5 mM Tris-HCl, pH 6.8, 5% 2-mercaptoethanol, 2% SDS, 5% sucrose, 0.005% bromophenol blue) by vortexing for 15 min at 37 °C. The samples were separated by SDS-PAGE. The separated samples were transferred to PVDF membranes. Membranes were blocked by 5% non-fat milk in Tris-Buffered Saline, with 0.002% Tween-20. Mouse anti-sera against GFP (mFx75, Wako), rabbit anti-sera against IbpA (Eurofin), and rabbit anti-sera against FtsZ (a gift from Dr. Shinya Sugimoto at Jikei Medical University) were used as primary antibodies at a 1:10,000 dilution. HRP conjugated anti-mouse IgG (Sigma-Aldrich) and HRP conjugated anti-rabbit IgG (Sigma-Aldrich) were used as secondary antibodies. Blotted membranes were detected with an LAS 4000 mini imager (Fujifilm).

### Quantitative RT-PCR

Total mRNA was extracted using Tripure Isolation Reagent (Merck) and treated with recombinant DNase I (Takara). The treated RNA was purified with an RNeasy Mini kit (Qiagen). The samples were prepared using a Luna Universal One-Step RT-qPCR Kit (New England Biolabs). Quantitative RT-PCR was performed with an Mx3000P qPCR system (Agilent) and analyzed by the MxPro QPCR software (Agilent). The amount of target mRNA was normalized with the *ftsZ* mRNA by the ∆∆Ct method [38]. Primers used for PCR are listed in *SI appendix Table S1*.

### Microscopy

To observe the IbpA localization in cells, we used the *E. coli* BW25113 wild-type strain carrying pCA24N-*rhodanese* and the *dnaKJ* deletion strain. Each strain carrying pBAD30-*ibpA* 5’ UTR-*ibpA-gfp* was grown to an OD_660_ of ~0.4 at 37 °C in LB medium. Cells were observed with an inverted microscope IX71 (Olympus) and a mercury lamp with a GFP filter. Fluorescent images were recorded with an iXon DV897 electron multiplying CCD camera (Andor) and the Andor SOLIS software (Andor).

### Protein purification

To purify IbpA, we used the BL21 (DE3) strain carrying pCA24N-*ibpA* or pCA24N-*ibpA*_AEA. Cells were grown in LB media at 37 °C to an OD_660_ of 1.0, and IbpA production was induced with 1 mM IPTG for 2 h. The cells overexpressing IbpA were lysed by sonication (Branson) in buffer A (50 mM Tris-HCl, pH 7.4, 10% glycerol, 1 mM dithiothreitol, 100 mM KCl). After the sonication, we followed the established methods for the purification of the wild-type IbpA [14] and the AEA mutant [31].

### Sucrose density gradient assay

The purified IbpA (10 µM) was incubated for 30 min at 48 °C in buffer B (50 mM Tris-HCl, pH 7.4, 150 mM KCl, 20 mM magnesium acetate, 5 mM dithiothreitol). After the incubation, the samples were applied onto an 11 ml gradient of 10-50% (w/v) sucrose in buffer B and centrifuged, using a Beckman SW41Ti rotor at 35,000 rpm at 4 °C for 80 min. The samples were collected as 20 separate fractions, using a fractionator (BioComp). The fractions were separated by SDS-PAGE and visualized by Coomassie Brilliant Blue staining.

### Growth assay

The *E. coli* wild-type strain, the *ibpAB* deleted strain, and the wild-type strain harboring pCA24N-*ibpA* or pCA24N-*gfp* were precultured at 30 °C for 16 h in LB medium. The precultured cells were incubated at 37 °C with shaking at 70 rpm, using a TVS062CA incubator (Advantec).

### Statistical Analysis

Student’s *t* test was used for calculating statistical significance, with a two-tailed distribution with unequal variance. All experiments represent a minimum of three independent experiments, with the bars showing the mean values ± SD.

### Data Availability Statement

Data in this manuscript have been uploaded on the Mendeley Dataset public repository (http://dx.doi.org/10.17632/jfnjyyyrfx.1).

## Acknowledgments

We thank Tatsuya Niwa for valuable discussions, Eri Uemura for technical support, Shinya Sugimoto for the anti-FtsZ antibody, the Bio-support Center at Tokyo Tech for DNA sequencing, and The National BioResource Project, *E. coli* (National Institute of Genetics, Japan) for providing the ASKA library clones and Keio collection strains. This work was supported by a MEXT Grant-in-Aid for Scientific Research (Grant No. 26116002 to HT).

## Supporting Information

**S1 Fig.**
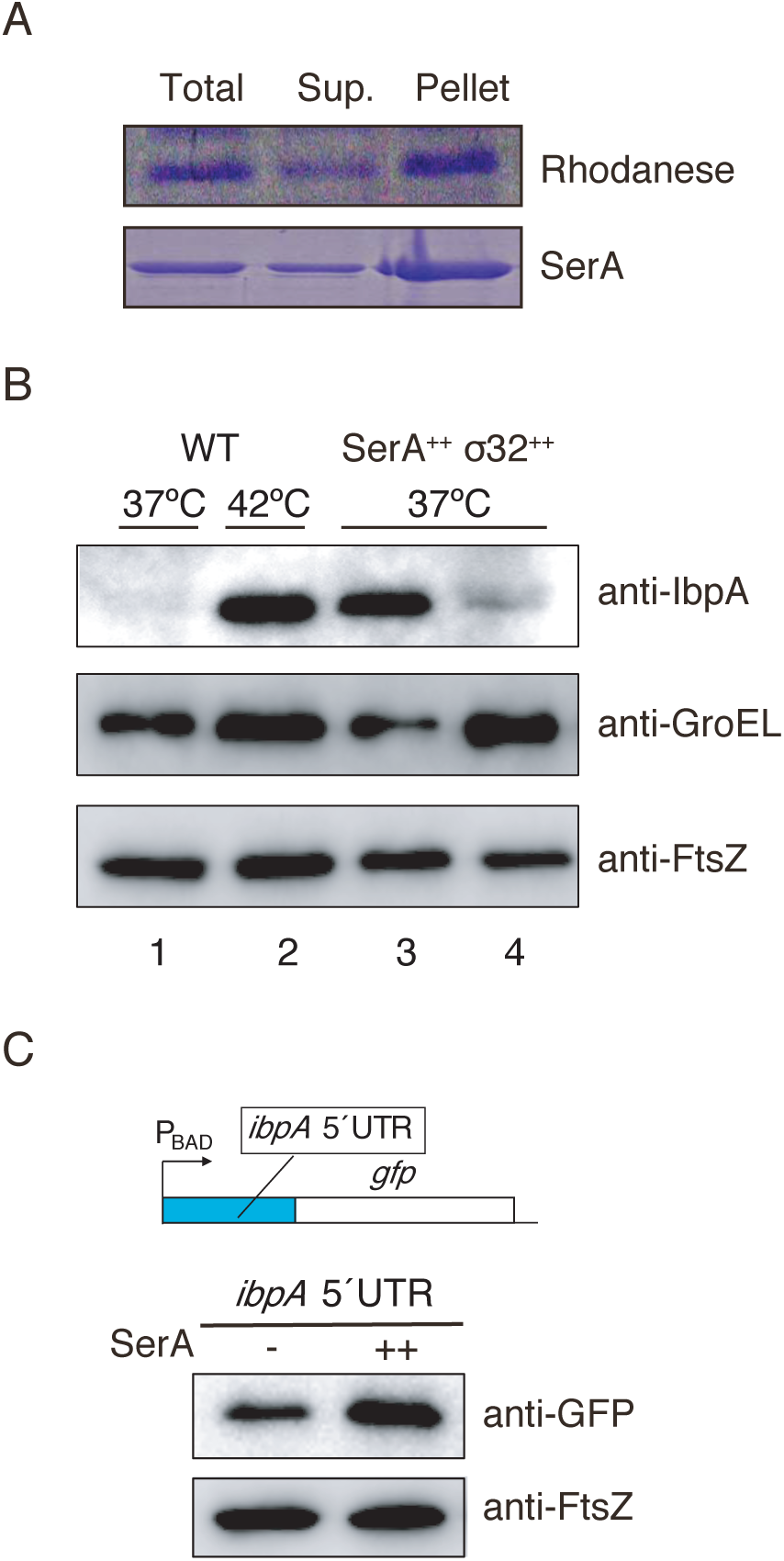
Overexpression of aggregation-prone proteins, rhodanese and SerA, in *E. coli*. (*A*) Aggregation formation of rhodanese and SerA in *E. coli* wild type upon the induction. The total, supernatant (*Sup*.) and pellet fractions of lysates from *E. coli* expressing rhodanese or SerA were analyzed by SDS-PAGE. The gel was stained with Coomassie blue. (*B*) Western blotting to evaluate the endogenous IbpA expression in *E. coli* wild-type under various conditions. 37 °C, *E. coli* grown at 37 °C; 42 °C, *E. coli* treated at 42 °C for 10 min. WT, *E. coli* wild-type; SerA_++_, *E. coli* wild-type expressing SerA; σ_32++_, *E. coli* wild-type expressing σ^32^. Expressions of GroEL and FtsZ were examined as controls for a representative Hsp and a constitutively expressed protein, respectively. Western blotting using anti-IbpA, anti-GroEL, and anti-FtsZ antibodies is shown. (*C*) Evaluation of the *ibpA* translation initiation by the GFP reporter used in Fig. 1*C*. *E. coli* lysates from wild type with (++) or without (–) the SerA induction by 0.1 mM IPTG for 1 h were analyzed. Western blotting using anti-GFP and anti-FtsZ antibodies is shown.

**S2 Fig.**
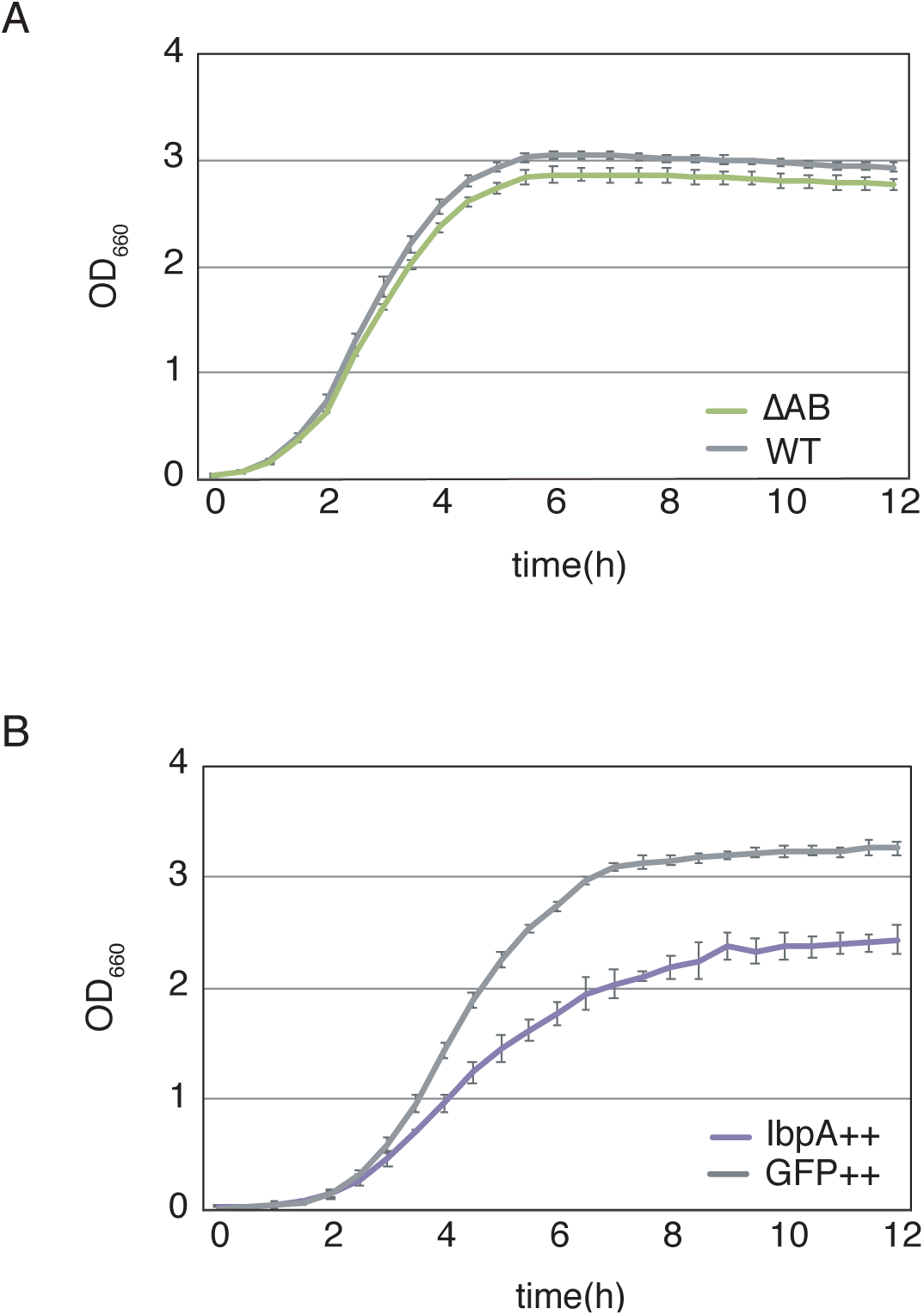
Cell growth of the *E. coli* strains. (*A*) Growth of *E. coli* wild-type (*WT*) and *ibpAB* deleted strain (∆*AB*). (*B*) Growth of *E. coli* wild-type strain overexpressing IbpA or GFP. Protein expression in the cultures harboring the plasmids was induced with 0.1 mM IPTG after 2 h from starting culture. Cells were grown at 37 ºC in LB medium in both experiments. The OD_660_ was measured every 30 min.

**S3 Fig.**
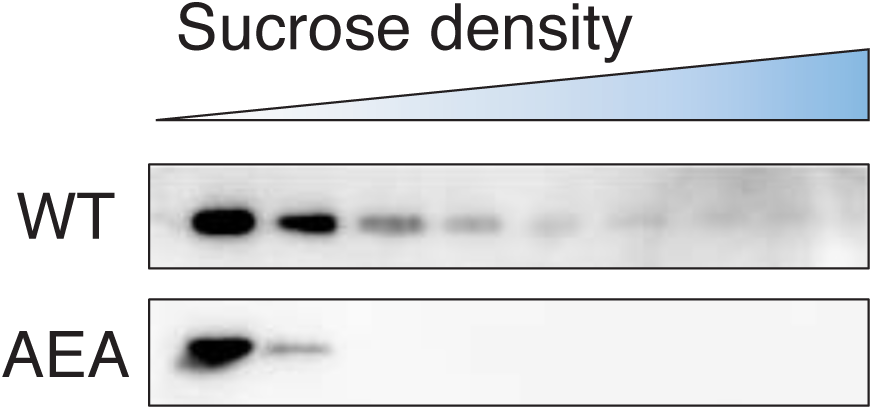
IEI motif in the C-terminal domain of IbpA is important for the IbpA oligomerization. Sucrose density gradient analysis of purified IbpA. The proteins were detected with western blotting using anti-IbpA antibody. *WT*, IbpA wild type; *AEA*, IbpA AEA mutant.

**S4 Fig.**
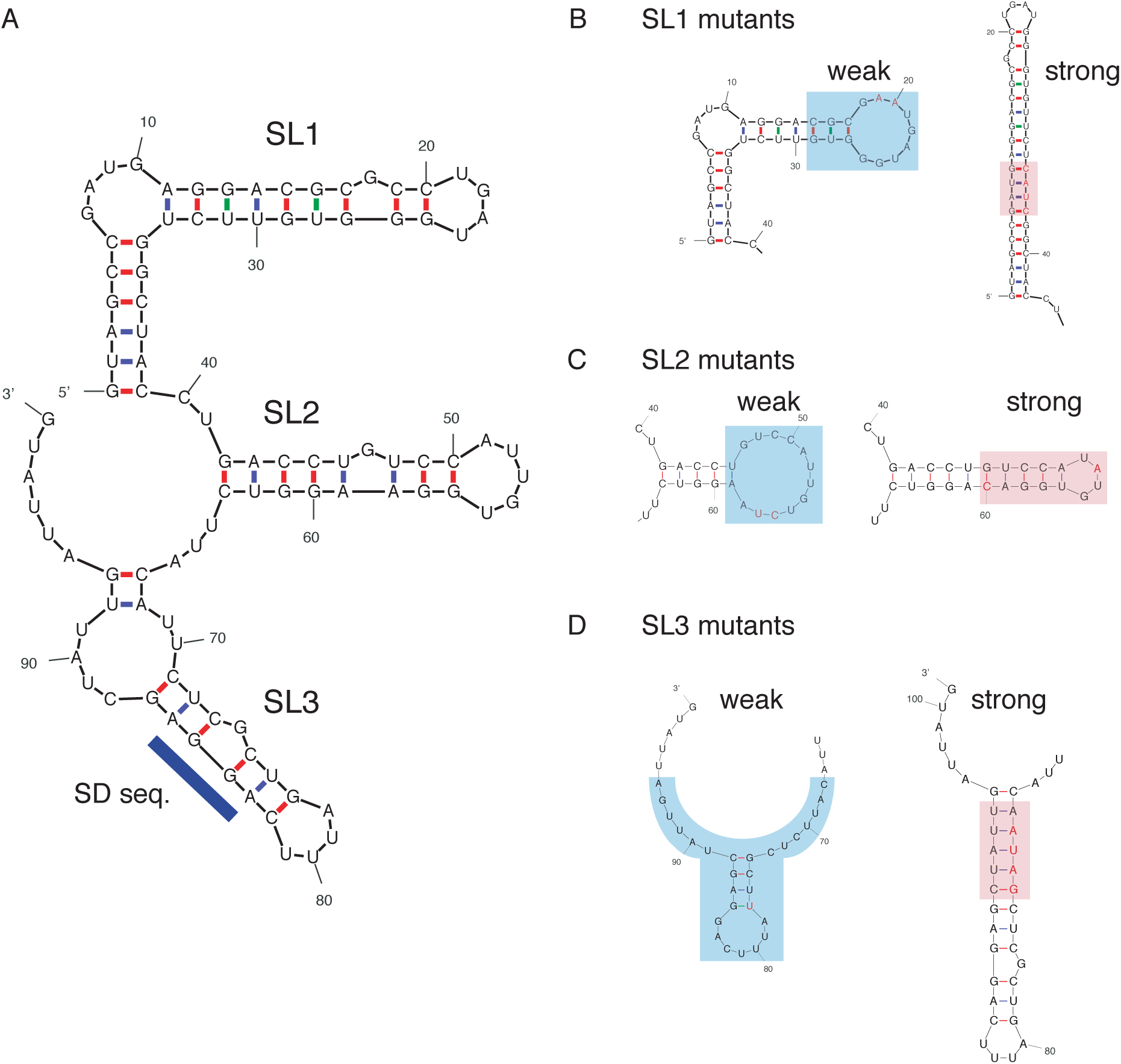
Secondary structure prediction of *ibpA* 5’ UTR mutants. The secondary structure prediction of *ibpA* 5’ UTR wild type (*A*) and mutants of stem loop 1-3 (SL1-3) (*B-D*). Color boxes represent the regions that were mutated to change the stability of stem loops. Blue boxes: weakened regions, Red boxes: strengthened regions.

**S1 Table.**
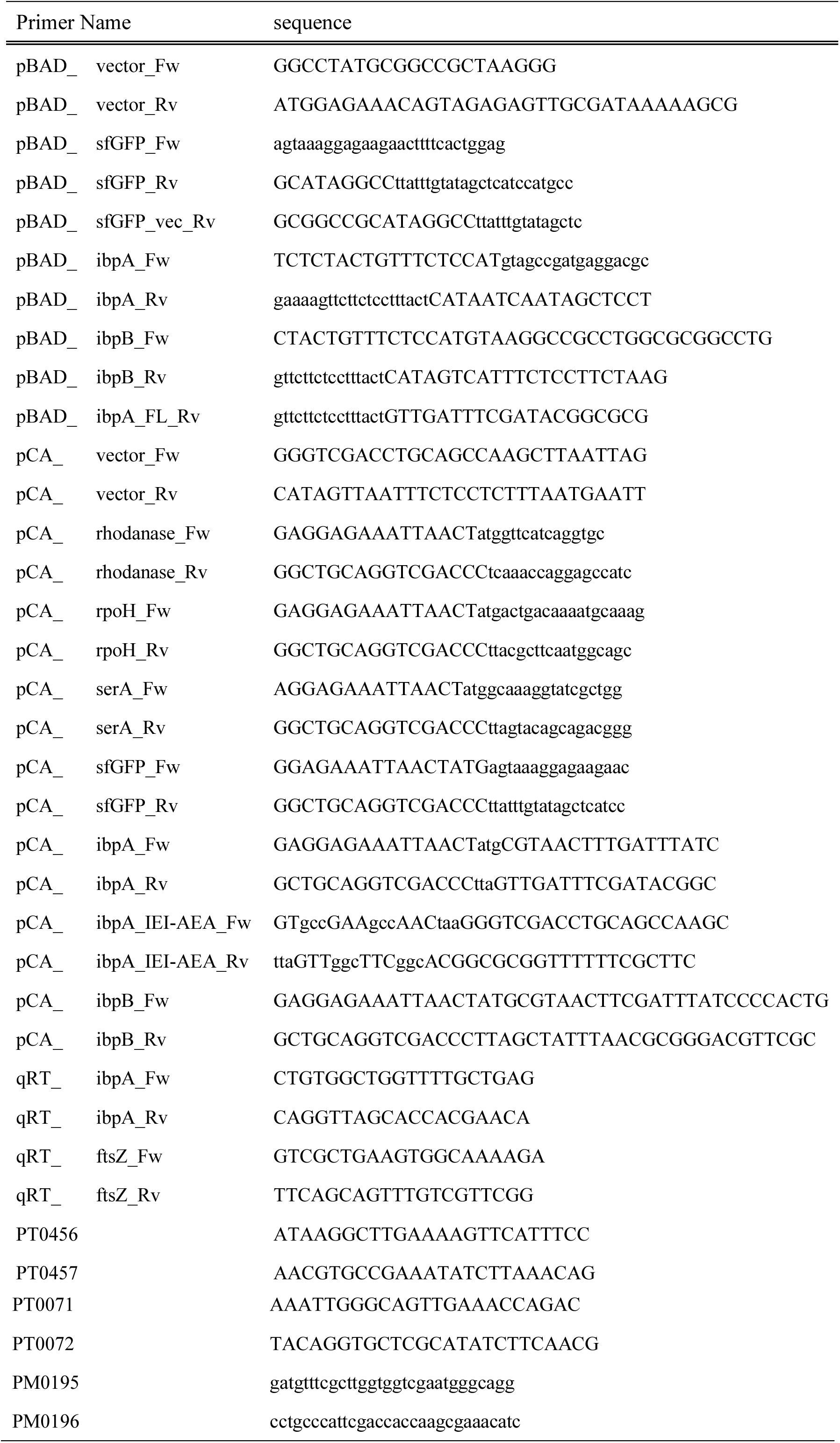
Primers used in this study.

